# A deoxynucleoside triphosphate triphosphohydrolase promotes cell cycle progression in *Caulobacter crescentus*

**DOI:** 10.1101/2024.04.25.591158

**Authors:** Chandler N. Hellenbrand, David M. Stevenson, Katarzyna A. Gromek, Daniel Amador-Noguez, David M. Hershey

## Abstract

Intracellular pools of deoxynucleoside triphosphates (dNTPs) are strictly maintained throughout the cell cycle to ensure accurate and efficient DNA replication. DNA synthesis requires an abundance of dNTPs, but elevated dNTP concentrations in nonreplicating cells delay entry into S phase. Enzymes known as deoxyguanosine triphosphate triphosphohydrolases (Dgts) hydrolyze dNTPs into deoxynucleosides and triphosphates, and we propose that Dgts restrict dNTP concentrations to promote the G1 to S phase transition. We characterized a Dgt from the bacterium *Caulobacter crescentus* termed *flagellar signaling suppressor C* (*fssC*) to clarify the role of Dgts in cell cycle regulation. Deleting *fssC* increases dNTP levels and extends the G1 phase of the cell cycle. We determined that the segregation and duplication of the origin of replication (*oriC*) is delayed in Δ*fssC*, but the rate of replication elongation is unchanged. We conclude that dNTP hydrolysis by FssC promotes the initiation of DNA replication through a novel nucleotide signaling pathway. This work further establishes Dgts as important regulators of the G1 to S phase transition, and the high conservation of Dgts across all domains of life implies that Dgt-dependent cell cycle control may be widespread in both prokaryotic and eukaryotic organisms.

**Importance:** Cells must faithfully replicate their genetic material in order to proliferate. Studying the regulatory pathways that determine when a cell initiates DNA replication is important for understanding fundamental biological processes, and it can also improve the strategies used to treat diseases that affect the cell cycle. Here, we describe a nucleotide signaling pathway that regulates when cells will begin DNA replication. We show that this pathway promotes the transition from the G1 to the S phase of the cell cycle in the bacterium *Caulobacter crescentus* and propose that this pathway is prevalent in all domains of life.

## Introduction

All cells proliferate through a highly ordered sequence of events known as the cell cycle. Precise coordination of the cell cycle is critical for survival, as improper control can lead to genome instability or cell death. For instance, the intracellular pools of deoxynucleoside triphosphates (dNTPs) must be strictly regulated throughout the cell cycle. As the precursors of DNA, physiological dNTP levels are crucial for accurate and efficient DNA replication(1). Perturbed dNTP levels can decrease polymerase fidelity, cause DNA damage, and stall replication forks(2–4).

The regulation of dNTP levels is coordinated with DNA synthesis(5). dNTP concentrations increase after the initiation of DNA replication to provide substrates for DNA polymerase. Ribonucleotide reductase (RNR) increases dNTP levels during DNA replication by synthesizing dNTPs from ribonucleoside triphosphates (rNTPs)(6–8). Its activity is upregulated by a variety of mechanisms after a cell enters S phase, and it is downregulated outside of S phase to reduce dNTP levels in nonreplicating cells(6, 9). Another family of enzymes known as deoxyguanosine triphosphate triphosphohydrolases (Dgts) helps regulate intracellular dNTP levels by hydrolyzing dNTPs into deoxynucleosides and triphosphates(5, 10). Dgts are present in all domains of life, but their physiological purpose remains less defined.

Few Dgts have been characterized, and differences in their catalytic mechanisms have led to nebulous conclusions about their functions(11–17). Dgts belong to a larger group of enzymes called the HD hydrolase superfamily(18). These enzymes harbor an HD motif that coordinates a divalent cation necessary for catalysis. All Dgts hydrolyze dNTPs through the same mechanism, but individual enzymes display a variety of substrate preferences. Dgts also vary in mechanisms of activation. Some enzymes require the binding of dNTPs at allosteric sites to activate dNTP hydrolysis, but the need for allosteric activation varies among different enzymes and depends on the identity of the cation present in the active site(16, 17, 19).

The mammalian Dgt, SAMHD1, reduces dNTP concentrations outside of S phase, and some have proposed that these enzymes restrict dNTPs as an antiviral strategy(10, 20, 21). Anti-viral roles for Dgts were originally predicted after T7 phage was found to encode an inhibitor of the *Escherichia coli* Dgt(22). Since then, many bacterial Dgts have been shown to increase phage resistance by limiting dNTPs and preventing the replication of viral genomes. SAMHD1 has also been identified as an HIV-1 restriction factor in human cells and is counteracted by the lentivirus auxiliary protein Vpx. (21, 23, 24). However, not all Dgts influence a host’s sensitivity to viral infection, and it is predicted that they have other physiological roles(20).

Elevated dNTP concentrations delay entry into S phase in eukaryotes, indicating that Dgts may have a role in cell cycle regulation. The depletion of SAMHD1 in human cells elevates dNTP pools and increases the steady-state proportion of cells in G1 phase(10). Deleting a Dgt in the protozoan *Trypanosoma brucei* yields a similar increase in the proportion of G1 cells(12). Overexpressing a constitutively activated RNR also extends G1 phase in *Saccharomyces cerevisiae* by increasing dNTP levels(25). These observations are counter intuitive given that dNTPs are programmed to increase during DNA synthesis, but they suggest the presence of an undefined regulatory mechanism through which high dNTP concentrations block the G1-S phase transition. We predict that Dgts maintain dNTP concentrations at a basal level in nonreplicating cells to promote the transition into S phase.

We have identified a Dgt from *Caulobacter crescentus* that establishes these enzymes as cell cycle regulators in bacteria. *C. crescentus* is a dimorphic bacterium that serves as an excellent model for studying the cell cycle (Fig. 1A)(26). There are two distinct *C. crescentus* cell types: motile swarmer cells and sessile stalked cells. Swarmer cells are incapable of initiating DNA replication; they must differentiate into stalked cells before they can enter S phase. Division in *C. crescentus* is asymmetric and yields one cell of each type. The stalked cell can immediately reenter S phase, but the swarmer cell will return to G1 phase and repeat the cycle(27).

**Figure 1:**
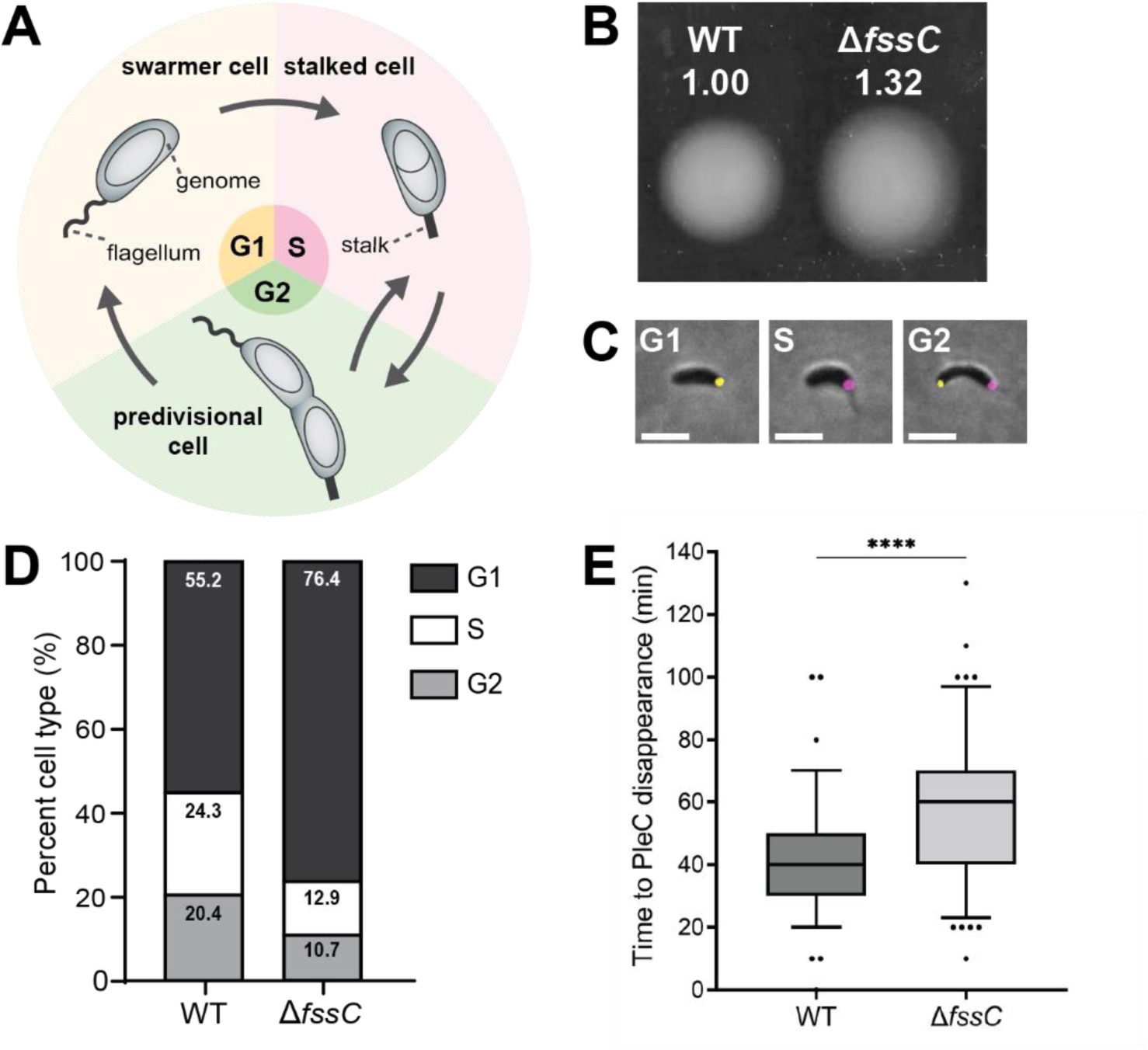
*fssC* promotes the swarmer-stalked transition. A) The dimorphic life cycle of *C. crescentus*. Motile swarmer cells are arrested in the G1 phase of the cell cycle and must differentiate into sessile stalked cells before entering S phase. B) The relative areas of WT and Δ*fssC* in a soft agar assay are shown. The Δ*fssC* mutant spreads 30% farther than WT through semi-solid medium. C) Example micrographs showing PleC-Venus (yellow) and DivJ-mKate (magenta) localization throughout the cell cycle. Scale bar is 2 µm. D) Measuring the localization of PleC-Venus and DivJ-mKate with fluorescent microscopy differentiates the phases of the *C. crescentus* cell. The Δ*fssC* mutant has a higher percentage (76.40%) of swarmer (G1) cells compared to WT (55.19%, *P =* 0.0067). Images were collected from unsynchronized CB15 populations in early exponential phase. Each bar represents n > 1000 cells collected over 3 biological replicates. E) Δ*fssC* has on average a 15.5-minute delay in the disappearance of its PleC-Venus foci. Box and whisker plots show the 5-95 percentile. Data was compiled over n=96 WT and n=105 Δ*fssC* unsynchronized cells. \*\*\*\**P* < 0.0001.

We identified the *C. crescentus* Dgt in a genetic screen designed to identify surface sensing genes(28). Swarmer cells use their flagellum to physically sense solid surfaces and activate signaling pathways that lead to surface attachment(28–30). Our group identified a panel of genes that are required to activate a surface response when the flagellum is disrupted. These *flagellar signaling suppressor* (*fss*) genes are predicted to activate surface adhesion downstream of surface sensing. We identified the *C. crescentus* Dgt as a putative surface sensing gene and named it *fssC* (Fig. S1).

This study aims to characterize the dNTP hydrolysis activity of FssC and its impact on cell cycle progression. Deleting *fssC* increases intracellular dNTP levels and delays entry into the stalked (S) phase of the cell cycle. We show that Δ*fssC* mutants have a delay in the segregation and duplication of the origin of replication (*oriC*) and conclude that elevated dNTPs inhibit the initiation of DNA replication. This study shows that Dgt-dependent cell cycle regulation is not restricted to eukaryotes and demonstrates that Dgts have important physiological roles beyond viral defense. We believe that Dgts regulate the cell cycle across all domains of life and propose that these enzymes are central to a novel dNTP signaling pathway that promotes the initiation of DNA replication.

## Results

### fssC promotes the swarmer (G1)-stalked (S) transition

1. *C. crescentus* cells migrate in semi-solid medium by using their flagella to chemotax through the agar matrix. Deleting *fssC* causes the cells to migrate 30% father than the WT strain (Fig. 1B). Given that *C. crescentus* is only motile during the swarmer phase of the cell cycle, this hyper-spreading phenotype can be indicative of a delay in the swarmer-stalked transition(31). We developed a fluorescence microscopy-based tool to quantify the proportion of swarmer cells in WT and Δ*fssC* populations. The histidine kinases PleC and DivJ were each fused to different colored fluorescent tags at their native loci. PleC localizes to the flagellar pole of swarmer cells and was fused to the yellow fluorescent protein Venus. DivJ localizes to the stalked pole of stalked cells and was fused to the red fluorescent protein mKate(32). These reporters allow for the visualization of each stage of the *C. crescentus* cell cycle (Fig. 1C). Swarmer cells are identified by a single PleC-Venus focus, stalked cells by a single DivJ-mKate focus, and predivisional cells by the presence of PleC and DivJ foci at opposite poles. The *pleC-venus* and *divJ-mKate* alleles did not substantially alter the motility phenotypes of WT or Δ*fssC* (Fig. S2). We analyzed unsynchronized populations of WT and Δ*fssC* with the *pleC-venus divJ-mkate* background and binned individual cells based on their PleC and DivJ localization. The Δ*fssC* mutant had a significantly higher proportion of swarmer cells (76.40%) compared to WT (55.19%), suggesting that this strain has an elongated swarmer phase (Fig. 1D).

Live cell microscopy was performed on unsynchronized cells to directly measure the duration of G1 phase in the Δ*fssC* mutant (Fig. 2E). WT and Δ*fssC* strains harboring *pleC-venus* were immobilized on agarose pads, and individual cells were imaged over a three-hour time-lapse experiment. The time required for PleC-Venus to delocalize in newly divided cells was recorded. PleC-Venus foci delocalize on average 15.5 minutes later in Δ*fssC* cells than in the WT background, confirming that *fssC* promotes the swarmer-stalked transition.

**Figure 2:**
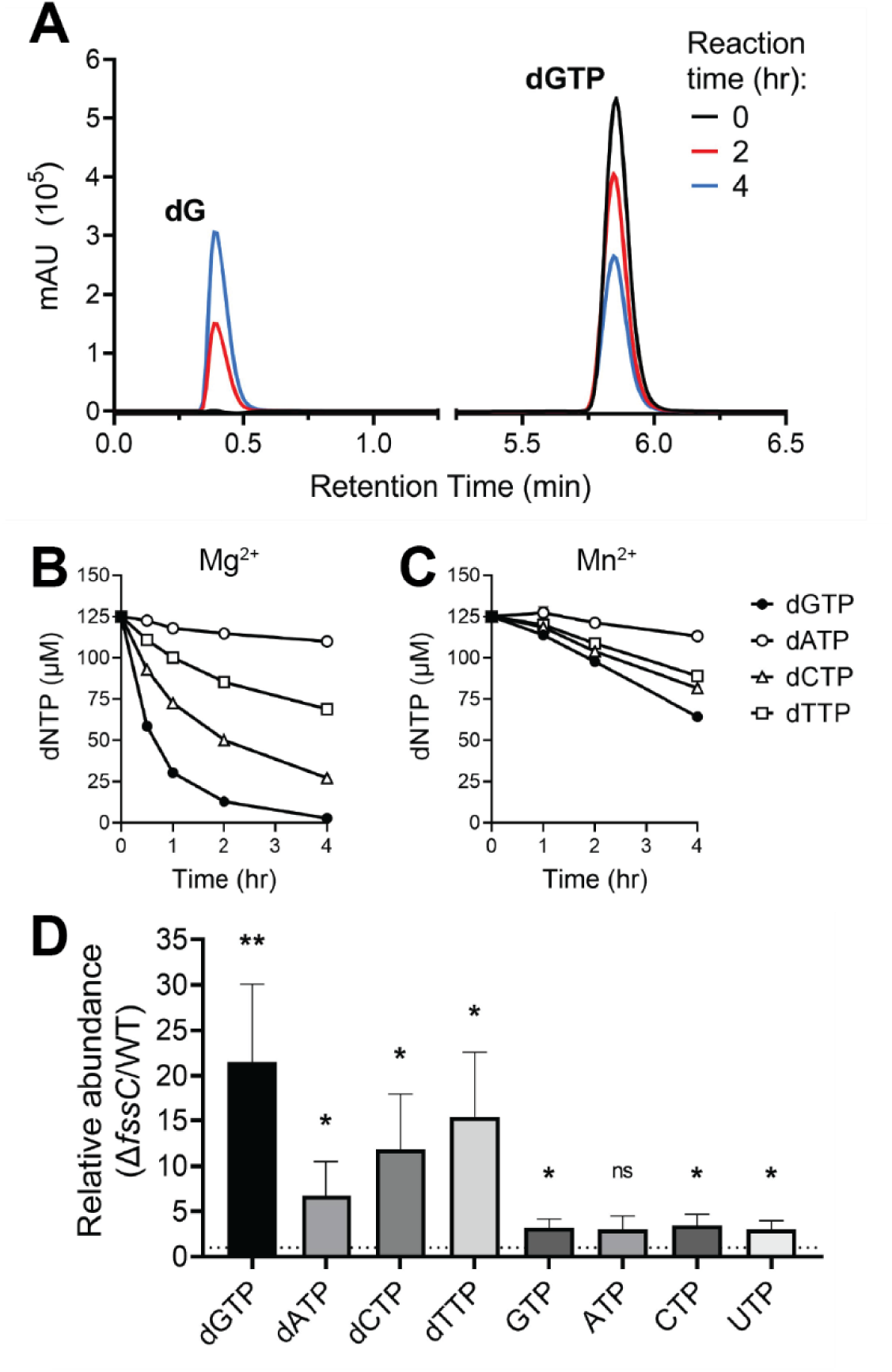
*fssC* encodes a deoxyguanosine triphosphate triphosphohydrolase (Dgt). A) dNTP hydrolysis was analyzed by anion exchange chromatography. Purified FssC was incubated with dGTP for 0 hrs (black), 2 hrs (red), and 4 hrs (blue) in reaction buffer supplemented with Mn^2+^. The dGTP substrate elutes at 5.85 min, and the dG product elutes at 0.388 min. B-C) FssC hydrolyzes all 4 canonical dNTPs *in vitro* with a preference for dGTP (dGTP > dCTP > dTTP > dATP). FssC was incubated with the 4 dNTPs mixed together (125 µM each, 500 µM total) in buffer containing Mg^2+^ (B) or Mn^2+^ (C) as the divalent cation. Error bars represent the standard deviation of the mean for three replicates. D) The Δ*fssC* mutant has higher intracellular dNTP levels than WT. Nucleotides were extracted from cell cultures and quantified by LC/MS. dGTP levels are on average 20 times higher in Δ*fssC* compared to WT. dATP, dCTP, and dTTP are also elevated but to a lesser extent. rNTP abundance is only slightly increased (∼3x higher in Δ*fssC*). Error bars represent the standard deviation of the mean for 3 biological replicates. \**P* < 0.05, \*\**P* < 0.01.

### FssC hydrolyzes dNTPs in vitro

The *fssC* gene encodes a predicted Dgt. A select group of Dgt homologs have been characterized, and individual enzymes display a variety of substrate preferences and activation mechanisms. The Dgts from *Escherichia coli* and *Leeuwenhoekiella blandensis* display strict specificity for dGTP and do not require allosteric activation(11, 13). TT1383 from *Thermus thermophilus* and EF1143 from *Enterococcus faecalis* hydrolyze all four canonical dNTPs but require allosteric activation by specific dNTP substrates(14, 16, 17). TT1383 and EF1143 only require activation when reaction buffer is supplemented with Mg^2+^ as the divalent cation. Replacing Mg^2+^ with Mn^2+^ circumvents the requirement for allosteric activation(16, 19).

We purified recombinant FssC and incubated the protein with various nucleotide substrates. Hydrolysis was assessed with anion exchange chromatography (Fig. 2A). FssC hydrolyzed each of the four dNTPs (dGTP, dATP, dCTP, dTTP), as measured by a decrease in the concentration of the dNTP substrate. Activity assays were performed with both individual dNTPs (Fig. S3) and with combinations of dNTPs. Two different reaction buffers were used (Fig. 2B, C). The first buffer contained Mg^2+^ as the divalent cation, and the second contained Mn^2+^. FssC demonstrates a clear kinetic preference for dGTP in either condition. However, dNTP hydrolysis only occurs in the Mg^2+^ buffer when FssC is incubated with dATP and at least one other dNTP (Table S1, Fig. S3). These results indicate that FssC requires activation by dATP when Mg^2+^ serves as the catalytic metal ion.

We tested FssC’s activity with a panel of potential nucleotide substrates. Assays were performed in three conditions: buffer supplemented with Mn^2+^, Mg^2+^, or with Mg^2+^ and dATP (FssC activating conditions). We examined deoxynucleotides (dGDP, dGMP), ribonucleotides (GTP), and the signaling nucleotides c-di-GMP (cdG), pppGpp, ppGpp, and pGpp. Hydrolysis by FssC was not detected for any of these substrates (Fig. S4).

We constructed a catalytically inactive FssC mutant by mutating the HD motif that coordinates the active site cation. Both residues (H102 and D103) were substituted for alanine. The FssC H102A D103A variant was unable to hydrolyze dNTPs in either Mg^2+^ or Mn^2+^ buffer (Fig. S5A). We used the H102A D103A variant to test if FssC’s hydrolysis activity was required for the enzyme to stimulate cell cycle progression. Expressing *fssC* from its native promoter at an ectopic locus in the Δ*fssC* mutant restores the wild-type motility phenotype in soft-agar. The hyper-spreading phenotype persists when the inactive H102A D103A variant is expressed in the mutant cells (Fig. S5B). This demonstrates that FssC’s catalytic activity is necessary for its role in regulating the swarmer-stalked transition. Indeed, ectopically expressing the H102A D103A mutant does not decrease the percentage of swarmers in the Δ*fssC* strain, while expressing wild-type *fssC* does (Fig. S5C). We conclude that dNTP hydrolysis by FssC is required for *C. crescentus* to efficiently progress though the cell cycle.

### FssC restricts intracellular dNTP concentrations

We used targeted metabolomics to examine the role of *fssC* in maintaining intracellular dNTP concentrations. Nucleotides were extracted from WT and Δ*fssC* cultures and analyzed by LC/MS to determine their relative abundance. The Δ*fssC* mutant has significantly higher dNTP levels than WT (Fig. 2D), and the relative abundance closely mirrors the *in vitro* substrate preference of the FssC enzyme (dGTP>dCTP>dTTP>dATP). dGTP is the most elevated dNTP in the Δ*fssC* mutant with levels 20 times higher than those in WT. dTTP and dCTP are 10-15 times higher in Δ*fssC*, and dATP is five times higher. The levels of rNTPs were also two to three times higher in Δ*fssC*. While it is possible that the FssC enzyme is more promiscuous *in vivo* than the *in vitro* hydrolysis assays indicate, we favor the explanation the elevated dNTPs could alter the flux of nucleotide metabolism and lead to a slight increase in rNTPs that is not a direct result of FssC activity. Regardless, this targeted metabolomic approach confirms that FssC is required to maintain low dNTP concentrations in *C. crescentus* cells.

### FssC does not affect the elongation phase of DNA replication

Elevated or imbalanced dNTP levels can be detrimental to the rate and fidelity of DNA replication(1–4). We therefore predicted that elevated dNTP concentrations in the Δ*fssC* mutant were influencing the rate of DNA replication. High-throughput sequencing was used to measure the DNA replication rate in WT and Δ*fssC* cells(33). A synchronizable strain of *C. crescentus* (NA1000) was used for these experiments. Populations were synchronized by isolating swarmer cells from a density gradient, and genomic DNA was sequenced at various time points after the cells were re-introduced into growth medium. The relative read coverage was plotted as a function of chromosome position to identify the location of the replication forks (Fig. 3C). Replication rates were calculated by plotting fork positions over time (Fig. 3B).

**Figure 3:**
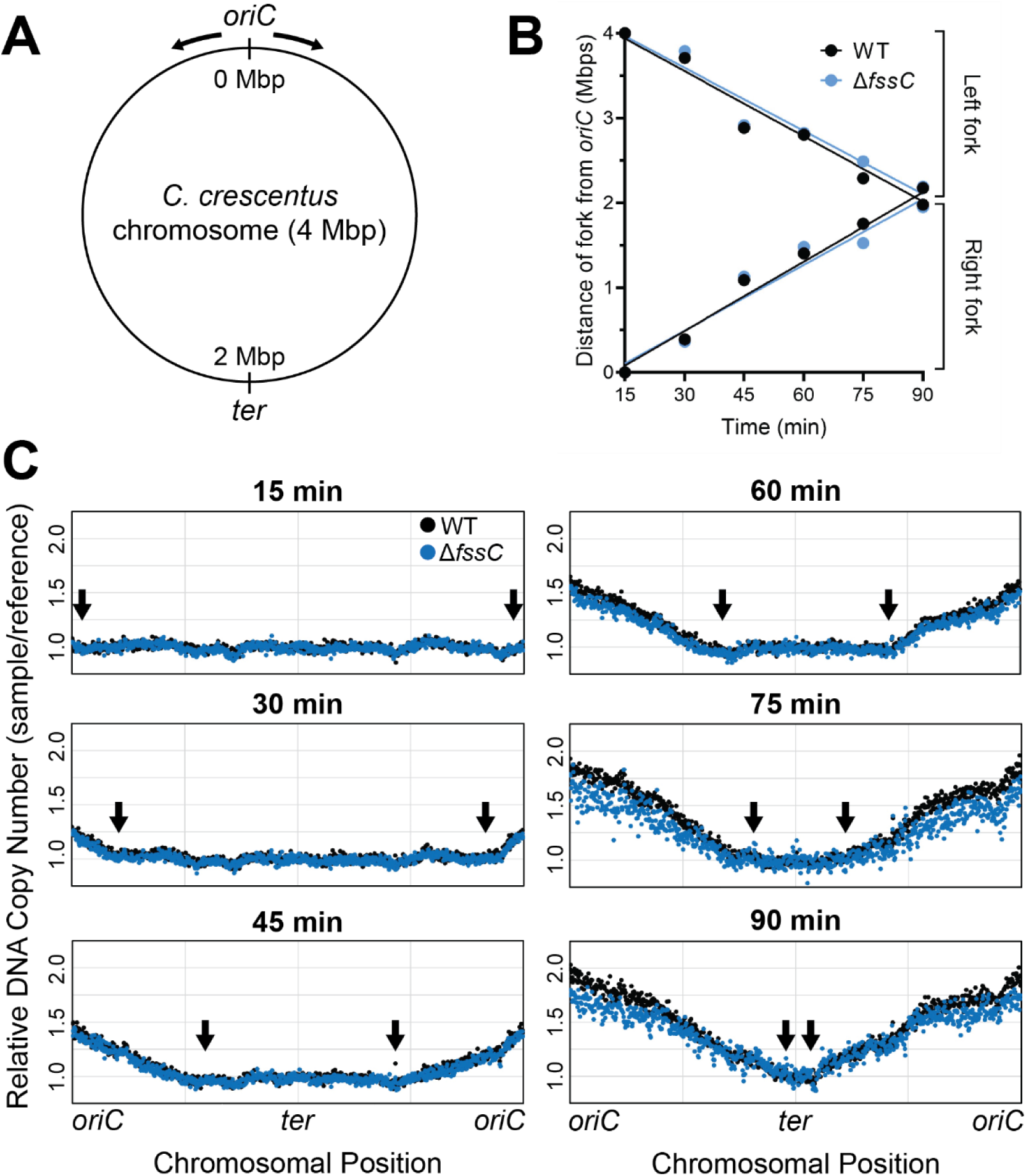
The Δ*fssC* mutant has a wild-type rate of DNA replication. A) The circular chromosome of *C. crescentus* is 4 Mbps in length. The origin of replication (*oriC*), terminus (*ter*) region, and the direction of the replication forks (black arrows) are shown. B) Positions of the right and left replication forks are plotted as a function of time for WT (black) and Δ*fssC* (blue). Line of best fit is shown for each fork. Slopes are not significantly different (*P* = 0.8547 for right forks, *P* = 0.7338 for left forks). C) Replication was monitored in synchronized NA1000 cells with a high-throughput sequencing approach. Read counts for each chromosomal position were normalized to t=0 to calculate relative copy number across the chromosome. Replication forks (black arrows) are at the interface between replicated and unreplicated DNA(34).

Replisomes on the left and right forks of the *C. crescentus* chromosome synthesize DNA at a rate of 428 ± 51 and 455 ± 39 bp/s, respectively. These rates are comparable to those found in *E. coli* and *B. subtilis*(34, 35). The rate of replication in the Δ*fssC* mutant is indistinguishable from the WT background. The left and right forks in Δ*fssC* move at a rate of 414 ± 50 and 432 ± 54 bp/s, respectively. This experiment was also performed with cells grown in M2X media (Fig. S6). We reasoned that cells in minimal media would grow slower and that any difference in replication between the two strains would be exacerbated. The results were comparable to the cells grown in PYE, confirming that the replication rates of WT and Δ*fssC* are identical. We conclude that elevated dNTP levels delay the swarmer-stalked transition through a mechanism independent of replication elongation.

### Segregation of the origin of replication (oriC) is delayed in ΔfssC

We next tested if the *ΔfssC* mutant has a delay in chromosome segregation. MipZ is a protein that associates with the centromere region near *oriC* on the *C. crescentus* chromosome(36). Fusing MipZ to a fluorescent Venus tag allows partitioning of the origin region to be tracked with live cell microscopy (Fig. 4A). We recorded the time required for newly divided swarmer cells to duplicate their Venus-MipZ foci as a measure of when the chromosomes begin to segregate. On average, Δ*fssC* cells duplicated their Venus-MipZ foci six minutes later than WT cells (Fig. 4B). A similar experiment was performed on NA1000 cells synchronized in the swarmer phase (Fig. S7). Venus-MipZ duplicates on average five minutes later in synchronized Δ*fssC*. These results indicate that the Δ*fssC* mutant has a delay in segregation of the origin region despite having a replication rate identical to WT.

**Figure 4:**
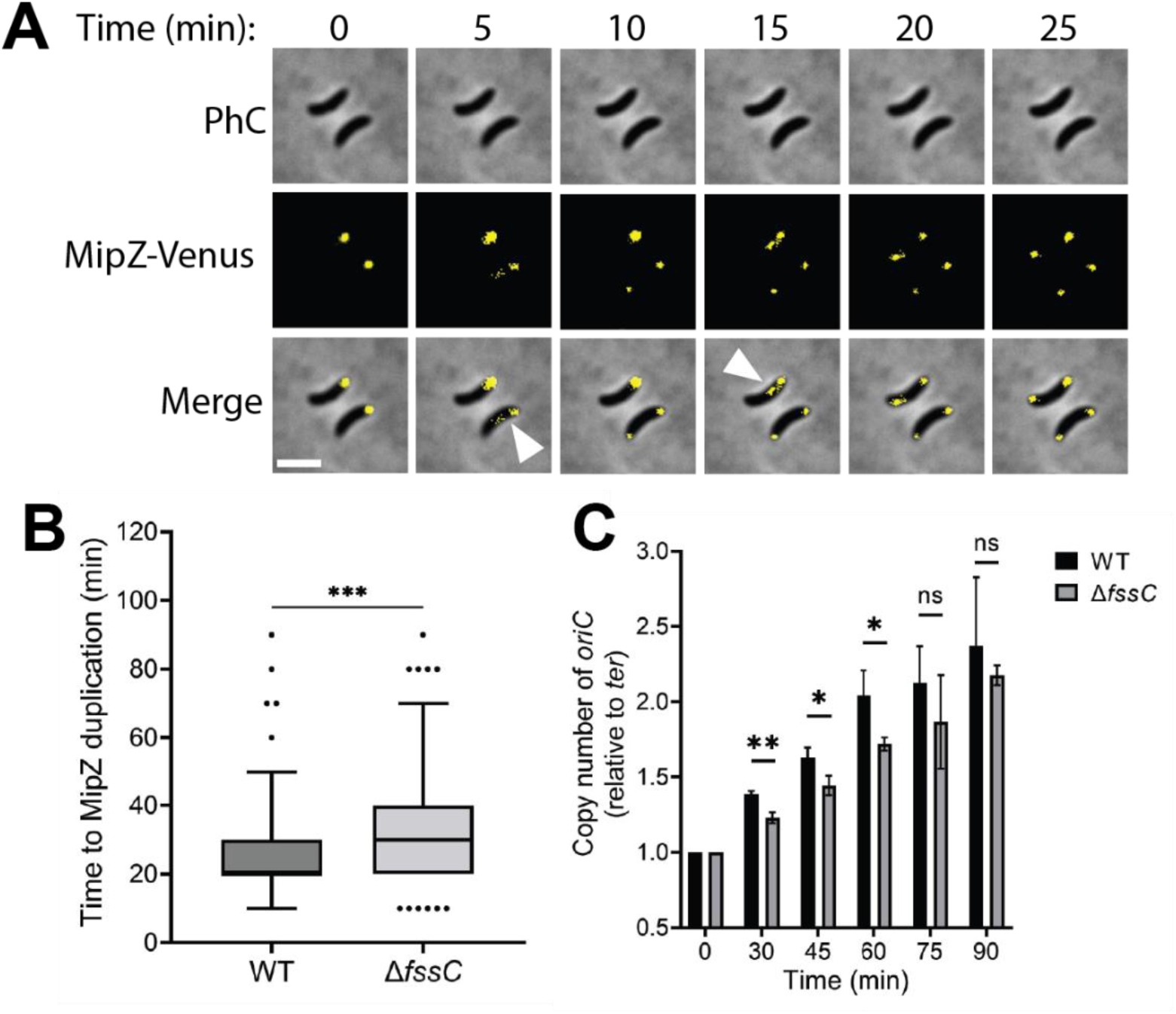
*fssC* promotes timely segregation and duplication of *oriC*. A) Representative micrographs showing the duplication of Venus-MipZ foci. Scale bar is 2 µm. B) Δ*fssC* has on average a 6-minute delay in MipZ duplication compared to WT. Box and whisker plots show the 5-95 percentile. Data is compiled from n=134 WT and n=182 Δ*fssC* unsynchronized cells. C) Relative copy number of *oriC* in synchronized populations over time was determined by qPCR. Primers were designed for *oriC* and the *ter* region (see Fig. 3A). The amount of *oriC* in each sample was normalized to the amount of *ter* and then to t=0. Error bars represent the standard deviation of the mean for 3 biological replicates. \**P* < 0.05, \*\**P* < 0.01, \*\*\**P* < 0.001.

### Initiation of DNA replication is delayed in ΔfssC

Given the identical replication elongation rates in WT and Δ*fssC*, we predicted that the delay in the segregation of the chromosomal origin reflected a delay in the initiation of DNA replication. A closer look at the high-throughput sequencing data (Fig. 3C and S6B) further supports this hypothesis. The copy number of *oriC* in the Δ*fssC* mutant is below that of WT at all timepoints.

We directly investigated the timing of replication initiation by measuring the relative copy number of *oriC* via quantitative PCR (qPCR). WT and Δ*fssC* cells were synchronized in the swarmer phase, and qPCR was performed on the *oriC* and the *ter* regions of the chromosome (Fig. 3A) to measure the *oriC*/*ter* ratio over time. When grown in PYE, the Δ*fssC* mutant has less *oriC* present than WT for up to 75 min post synchronization, at which point both strains have fully duplicated their origins and reached a copy number of 2N (Fig. 4C). qPCR was also performed on samples grown in M2X, yielding similar results (Fig. S6C). These data support the model that Δ*fssC* has a delay in the initiation of DNA replication and suggests that dNTP hydrolysis by FssC plays an important role in regulating entry into S phase.

## Discussion

dNTP levels are precisely regulated throughout the cell cycle to avoid DNA damage and promote efficient DNA replication(1–5). Over the last decade, it has become clear that Dgts play an important role in regulating dNTP levels(5, 10). Dgts reduce dNTP concentrations by hydrolyzing dNTPs into deoxynucleosides, but the physiological purpose of these enzymes remains debated.

We have characterized a Dgt in *C. crescentus* called *flagellar signaling suppressor C* that regulates the G1 to S phase transition of the cell cycle. *In vitro* characterization confirmed that the FssC enzyme has dNTP triphosphohydrolase activity. FssC has a kinetic preference for hydrolyzing dGTP but can hydrolyze all four canonical dNTPs (Fig. 2). FssC has a similar activation mechanism to two other characterized Dgts: TT1383 from *T. thermophilus* and EF1143 from *E. faecalis*(14, 16). All three enzymes require activation by dNTPs when reaction buffer is supplemented with Mg^2+^(14, 16, 17). We found that FssC requires dATP and at least one other dNTP to activate hydrolysis (Fig. S3, Table S1). Like TT1383 and EF1143, FssC does not require activation when supplemented with Mn^2+^. The dNTP hydrolysis activity of FssC is also relevant *in vivo*. We confirmed that dNTP levels are elevated in the Δ*fssC* mutant compared to WT with targeted metabolomics (Fig. 2D). All four canonical dNTPs are at least five times higher in Δ*fssC*, and dGTP was the most elevated with levels 20 times higher than WT.

We suspected *fssC* may have a role in controlling the cell cycle after examining motility in semi-solid agar (Fig. 1B). The Δ*fssC* mutant migrates 30% farther than WT, and we predicted that this result was caused by an elongated swarmer phase. This hypothesis was supported by fluorescent microscopy experiments that tracked the localization of PleC-Venus and DivJ-mKate (Fig. 1C, D). A delay in the delocalization of PleC-Venus from the swarmer pole confirmed that *fssC* is required for the timely transition from swarmer to stalked cell (Fig. 1E). This cell cycle phenotype is dependent on the dNTP hydrolysis activity of the FssC enzyme. The expression of a catalytically inactive *fssC* allele (H102A D103A) is unable to restore the WT phenotype in soft agar or the length of the swarmer phase (Fig. S5).

The Δ*fssC* mutant’s delay in the swarmer-stalked transition can be traced back to a delay in the initiation of DNA replication. Time-lapse microscopy of WT and Δ*fssC* strains harboring a fluorescent Venus-MipZ fusion showed that the Δ*fssC* mutant has a delay in segregation of the chromosomal origin of replication (Fig. 4B, S7). However, we found that WT and Δ*fssC* have identical rates of replication elongation (Fig. 3, S6). This led us to hypothesize that the deletion of *fssC* causes a delay in the initiation of DNA replication. We determined the relative copy number of *oriC* by performing qPCR on genomic DNA and found that the Δ*fssC* mutant had less *oriC* than WT (Fig. 4C, S6C). This result indicates that Δ*fssC* cells on average duplicate their *oriC* later than WT cells. We conclude that elevated dNTP levels in the Δ*fssC* mutant delay the initiation of DNA replication.

An analogous phenotype has previously been associated with elevated dNTP concentrations in eukaryotic systems. Removing Dgts from human fibroblasts and the protozoan parasite *T. brucei* increases dNTP levels and delays entry into S phase(10, 12). Inducing high dNTP concentrations by constitutively activating RNR has a similar effect on the cell cycle in *S. cerevisiae*(25). These findings are counter intuitive from a biochemical perspective. As substrates for DNA polymerase, elevated dNTPs are expected to promote DNA replication. The opposite has now been observed in two different domains of life. We propose that elevated dNTPs target the initiation of DNA replication through a novel nucleotide signaling pathway and that Dgts reduce dNTP levels for efficient progression of the cell cycle (Fig. 5).

**Figure 5:**
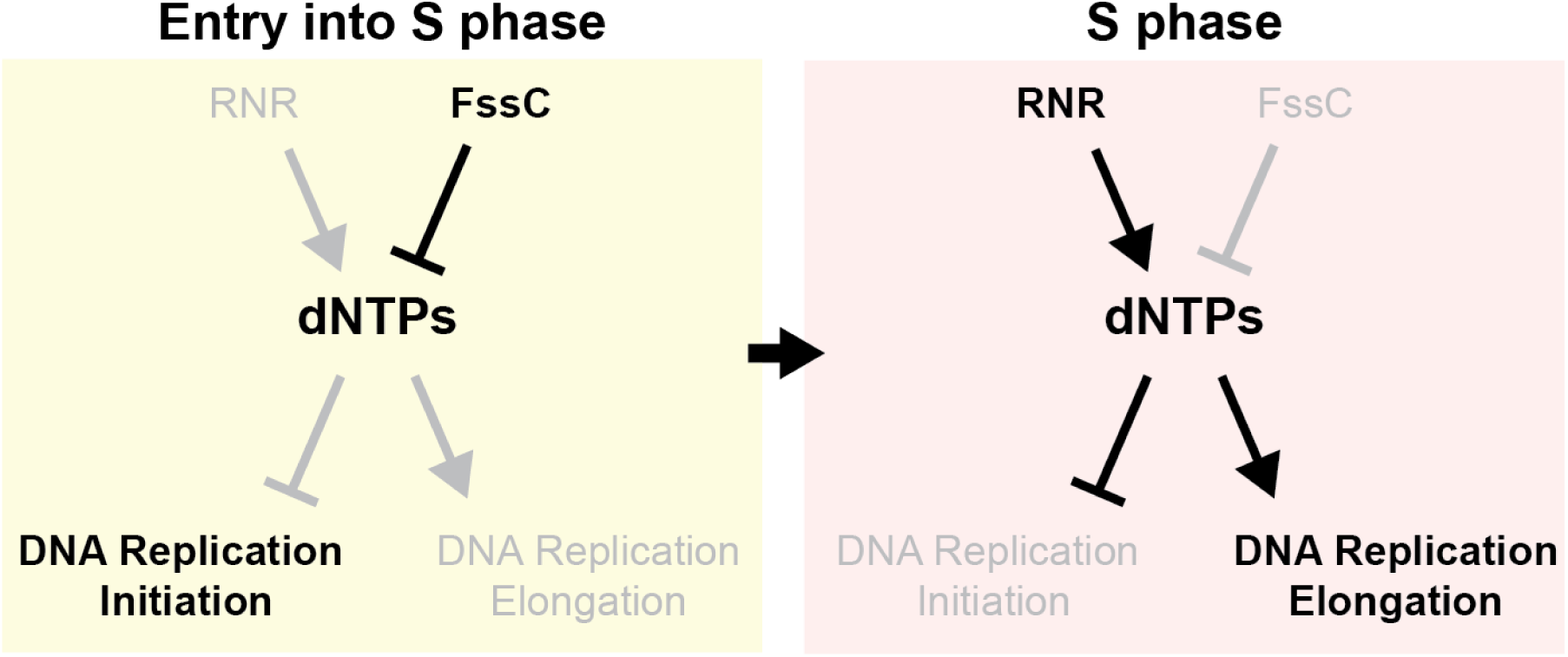
FssC promotes entry into S phase through a dNTP signaling pathway. Elevated dNTPs in G1 phase inhibit the initiation of DNA replication. Hydrolysis by FssC lowers dNTP levels to relieve this inhibition. Once in S phase, RNR is activated to increase dNTP levels and promote DNA synthesis.

The mechanism by which elevated dNTPs delay the initiation of DNA replication remains undefined. Some evidence suggests that high dNTPs delay the formation of the pre-initiation complex in yeasts(25), but bacteria do not have a pre-initiation complex. It is possible that elevated dNTPs have different targets in prokaryotic and eukaryotic organisms, but further studies are needed to determine the precise mechanism of this nucleotide signaling pathway.

Lowering dNTP concentrations can also affect the replication of invading viral genomes. Recent studies have proposed that Dgts hydrolyze dNTPs primarily as an antiviral strategy(20, 21). The depletion of dNTP pools can limit viral replication, and some bacterial Dgts are encoded next to other known phage defense genes(20). However, not all Dgts improve a host’s resistance to viral infection(20). For instance, we found that *fssC* does not influence the susceptibility of *C. crescentus* to ɸCbK infection (Fig. S9). We propose that increased viral defense is an indirect effect of dNTP hydrolysis, and that the primary role of Dgts is to promote entry into S phase.

Interestingly, we identified *fssC* in a suppressor screen that sought to discover new genes in the flagellum-mediated surface sensing pathway (Fig. S1)(28). Swarmer cells use both their flagella and type IV pili to physically sense surface contact and activate surface adhesion(29, 30, 37, 38). Previous studies have shown that the *C. crescentus* cell cycle is regulated by pilus-mediated surface contact and that physical obstruction of pilus retraction stimulates early entry into S phase(39, 40). The *fssC* signaling pathway may be an additional mechanism by which surface contact promotes chromosome replication in *C. crescentus.* Further experiments will be required to confirm that *fssC* is a true surface sensing gene, as the possibility that *fssC* functions as an independent regulator of the cell cycle cannot currently be ruled out.

We have shown that *fssC* regulates the G1 to S phase transition in *C. crescentus.* Elevated dNTP levels delay the initiation of DNA replication, and *fssC* promotes entry into S phase through dNTP hydrolysis (Fig. 5). Analogous phenotypes have been observed in eukaryotic organisms, and we predict that Dgt-dependent cell cycle regulation is widespread across the tree of life.

## Materials and Methods

### Bacterial strains, growth, and genetic manipulation

Strains used in this study are listed in Table 1. Plasmids (Table 2) were developed with PCR, restriction digestion, and Gibson assembly. Primer sequences are available upon request. *E. coli* was grown in LB medium at 37°C and supplemented with 50µg/mL kanamycin when necessary.

**Table 1:** Plasmids used in this study.

| Plasmid | Description | Antibiotic | Reference |
| --- | --- | --- | --- |
| pNPTS138 | Suicide plasmid for making unmarked deletions in *C. crescentus*; carries *sacB* for counter-selection | Km | M. R. Alley |
| pDH118 | pMT585, pMTLS4259, integrating vector for xylose inducible expression of C-terminally GFP tagged proteins in *Caulobacter* | Km | (44) |
| pDH804 | To delete *fssC*; Gibson cloning of fused upstream and downstream regions of CC_2008 | Km | This work |
| pDH1161 | To fuse fluorescent mVenus protein to C-terminus of PleC; Gibson cloning of *mvenus* fused between 3' end and downstream region of CC_2482 | Km | This work |
| pDH1209 | To fuse fluorescent mKate2 protein to C-terminus of DivJ; Gibson cloning of *mkate2* fused between 3' end and downstream region of CC_1063 | Km | This work |
| pDH430 | Modified pET28a plasmid that includes an 8xHis-SUMO tag upstream of multicloning site | Km | This work |
| pDH815 | To overexpress FssC in *E. coli*; 8xHis-SUMO-FssC cloned into pDH430. | Km | This work |
| pDH1230 | To overexpress FssC H102A D103A in *E. coli*; Quickchange mutagenesis of pDH815 | Km | This work |
| pDH1335 | To fuse fluorescent mVenus protein to N-terminus of MipZ; Gibson cloning of *mvenus* fused between upstream region and 5' end of CC_2165 | Km | This work |
| pDH1158 | To integrate *PfssC*-empty at *xyl* locus; Gibson cloning of CC_2008 promoter (88 nt upstream) | Km | This work |
| pDH1167 | To integrate *PfssC-fssC* at *xyl* locus; Gibson cloning of CC_2008 fused to CC-2008 promoter (88 nt upstream) | Km | This work |
| pDH1224 | To integrate *PfssC-fssC* H102A D103A at *xyl* locus; Quickchange mutagenesis of pDH1167 | Km | This work |
| pDH418 | To delete *flgH*; Gibson cloning of fused upstream and downstream regions of CC_2066 | Km | (45) |
| pDH805 | To delete *fliF*; Gibson cloning of fused upstream and downstream regions of CC_0905 | Km | (28) |

**Table 2:**
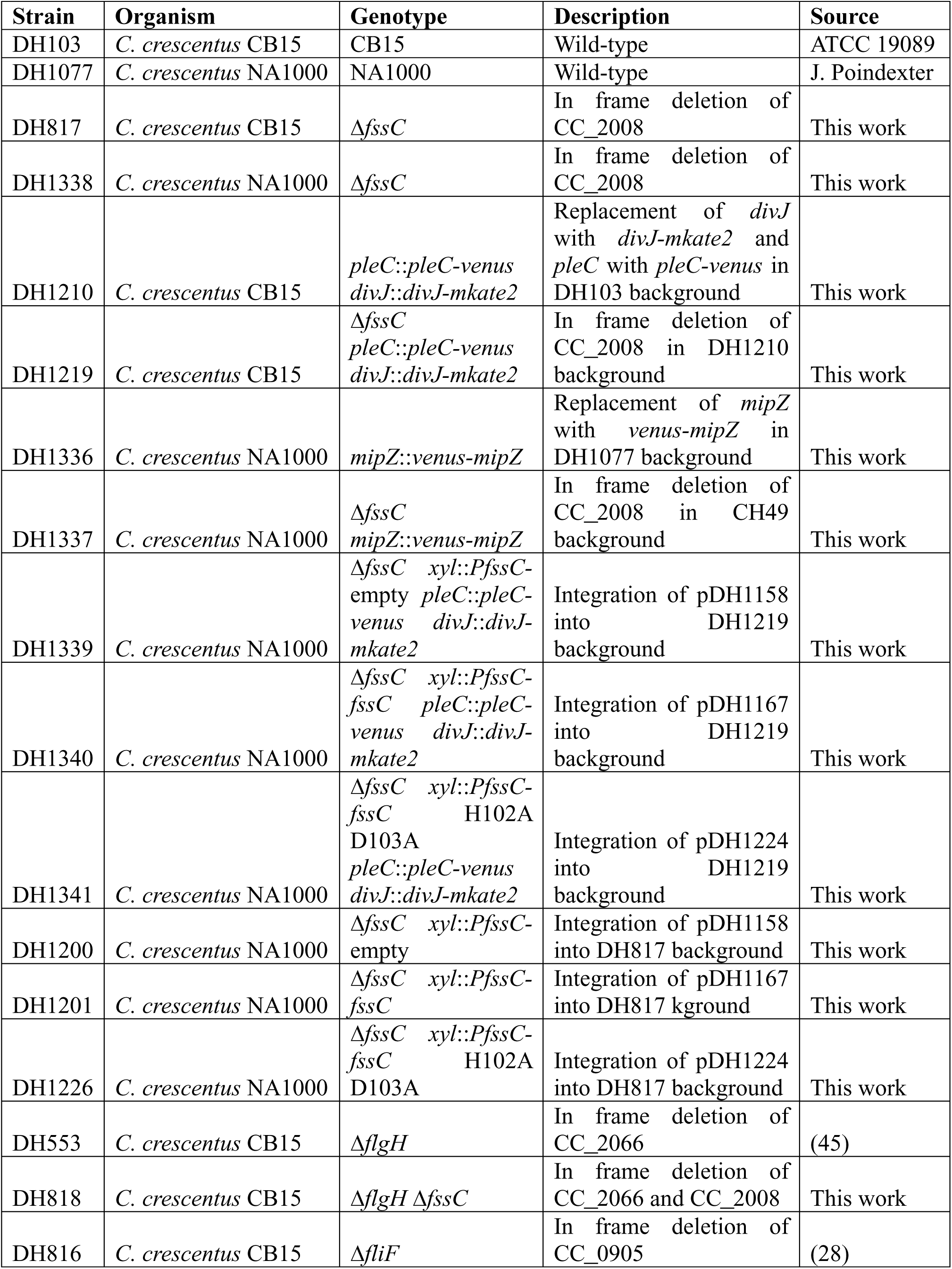
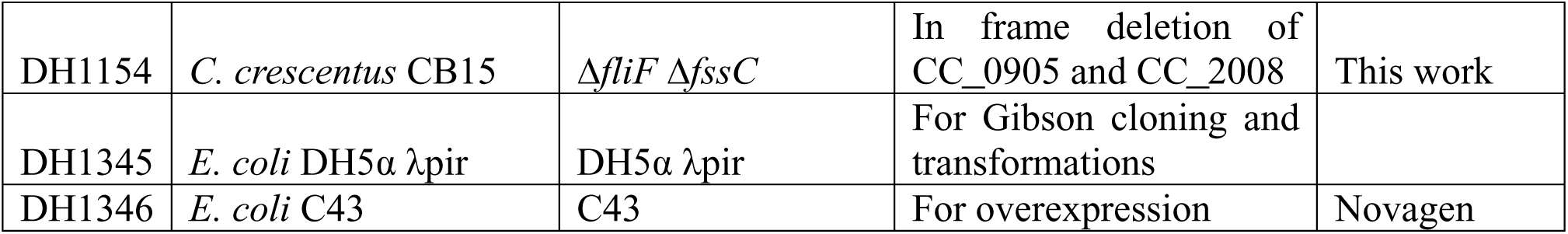
Strains used in this study.

*C. crescentus* was grown in PYE medium or M2 minimal media supplemented with 0.15% xylose (M2X) at 30°C. Liquid and solid PYE medium was supplemented with 5µg/mL and 25µg/mL kanamycin, respectively, when required. Plasmids were transformed into *C. crescentus* by electroporation. Gene deletions and insertions were constructed with a two-step approach using *sacB-*based counterselection.

### Soft agar motility assay

Strains were grown overnight in PYE and diluted to an OD_660_ of 0.5 before inoculating 2µL into PYE plates containing 0.3% agar. Plates were incubated at 30°C for 72 hrs and area of growth was measured.

### Determining cell cycle phenotypes of CB15 populations

Strains were grown overnight in PYE, then diluted to an OD_660_ of 0.05 and grown for 90 min. 2µL of cells were immobilized on a 1% agarose pad and imaged. Microscopy was performed using a Nikon Ti-E inverted microscope equipped with an Orca Fusion BT digital CMOS camera (Hamamatsu). Fluorescence images were collected using a Prior Lumen 200 metal halide light source and a YFP- and mCherry-specific filter set (Chroma). Image analysis was performed with MicrobeJ(41).

### Live cell imaging of PleC-Venus and Venus-MipZ in unsynchronized cells

Strains were grown overnight in PYE, then diluted to an OD_660_ of 0.05 and grown for 3 hrs. 2µL of cells were spotted onto a 1.5% agarose pad made with PYE and incubated at 30°C for 1 hr. Microscopy was performed with the same equipment described previously. Images were collected every 10 min for 3 hrs.

### NA1000 synchronization

Strains were grown overnight in PYE, diluted to an OD_660_ of 0.1 in M2X, and grown for 6-8 hrs. Cultures were diluted again into M2X and grown to an OD_660_ of 0.5-0.6. Cells were harvested by centrifugation and resuspended in chilled M2 salts and 1 volume of percoll. Swarmer cells were separated from stalked and predivisional cells by centrifugation at 15,000x*g* for 20 min at 4°C. The bottom swarmer band was collected and washed with M2 salts.

### Overexpression and purification of FssC

A pET28a vector encoding 8xHis-SUMO-FssC was transformed into *E. coli* strain C43. Transformants were grown overnight and diluted (1/100) into 1L of 2xYT media. Cultures were induced with 0.5mM IPTG at an OD_600_ of 0.35 and incubated for 4 hrs at 37°C. Cells were harvested by centrifugation and stored at -80°C.

Cell pellets were resuspended in 30mL lysis buffer (20mM Tris-HCl pH 7.4, 1M NaCl, 20mM imidazole, 1µM PMSF, and 10% glycerol) and passaged through a cell disruptor at 20,000psi until fully lysed. Lysates were centrifuged at 30,000x*g* for 20 min at 4°C. The supernatant was supplemented with 0.1% PEI pH 7.25 and centrifuged again at 50,000x*g* for 20 min at 4°C. 5mL of Ni-NTA resin was added to the supernatant, and the slurry was rocked for 1 hr at 4°C. The resin was washed with NWB (20mM Tris-HCl pH 7.4, 300mM NaCl, 10mM imidazole, 10% glycerol), and protein was eluted with NEB (20mM Tris-HCl pH 7.4, 800mM NaCl, 500mM imidazole, 10% glycerol). 6xHis-Ulp1 enzyme was added to the eluate and dialyzed against DC buffer (25mM Tris-HCl pH 7.4, 300mM NaCl, 10mM imidazole, 10% glycerol).

Cleavage reaction was transferred onto a column with 3mL Ni-NTA resin. The flowthrough was concentrated with an Amico Ultra 30,000 MWCO. Concentrated sample was further purified with size exclusion chromatography using an AKTA Pure (GE Healthcare) FPLC system with a HiPrep 26/60 Sephacrul S-200 column. Fractions containing FssC were pooled, concentrated, and stored at -80°C.

### dNTP hydrolysis assays

Assays were performed in 50mM Tris-HCl pH 8, 100mM NaCl, 0.4mM DTT, and either 5mM MnCl_2_ or MgCl_2_. All reactions contained 100µM purified FssC and were incubated at 30°C. 65mM EDTA was added to quench reactions. FssC was precipitated by 1 volume of chilled methanol. Samples were analyzed by anion exchange using a DNAPac PA-100 (4 X 50mm) column on a Shimadzu LC40 HPLC equipped with an SPD-M40 photodiode array detector. For reactions containing a single dNTP, the column was equilibrated with 25mM Tris-HCL pH 7.4 and 0.5mM EDTA (buffer A). Injected sample (20µL) was eluted with a 3 min isocratic phase of buffer A followed by a 10 min linear gradient of 0-500mM LiCl. For reactions containing dGMP and/or multiple nucleotides, the column was equilibrated with 2.5% acetonitrile, and injected samples (25µL) were eluted with a 25 min linear gradient of 0-175mM potassium phosphate pH 4.6. Absorbance was continuously monitored between 200 and 500nm. Nucleotides were quantified by peak integration at 260nm.

### Quantification of intracellular dNTPs

Nucleotides were extracted from cell cultures as described previously(42). Cultures were grown to an OD_660_ of 0.3-0.5 in M2X, and cells were harvested by vacuum filtration with a PTFE membrane (Satorius, SAT-11806-47-N). The membrane was submerged in chilled extraction solvent (50:50 (v/v) chloroform/water). Extracts were centrifuged to remove cell debris and the organic phase. The aqueous layer was stored at -80°C.

Samples were analyzed using an HPLC-tandem MS (HPLC-MS/MS) system consisting of a Vanquish UHPLC system linked to heated electrospray ionization (HESI) to a hybrid quadrupole high resolution mass spectrometer (Q-Exactive orbitrap, Thermo Scientific) operated in full-scan selected ion monitoring (MS-SIM) using negative mode to detect targeted metabolites. MS parameters included: a resolution of 70,000, an automatic gain control (AGC) of 1e6, spray voltage of 3.0kV, a maximum ion collection time of 40 ms, a capillary temperature of 35°C, and a scan range of 70–1000mz. LC was performed on an Aquity UPLC BEH C18 column (1.7μm, 2.1 × 100mm; Waters). 25µL of the sample was injected via an autosampler at 4°C. Total run time was 30 min with a flow rate of 0.2 mL/min, using Solvent A (97:3 (v/v) water/methanol, 10mM tributylamine (Sigma-Aldrich) pH∼8.2–8.5 adjusted with ∼9mM acetic acid) and 100% acetonitrile as Solvent B. The gradient was as follows: 95% A/5% B for 2.5 min, then a gradient of 90% A/10% B to 5% A/95% B over 14.5 min, then held for 2.5 min at 10% A/90% B. Finally, the gradient was returned to 95% A/5% B over 0.5 min and held for 5 min. HPLC eluate was sent to the MS for data collection from 3.3 to 18 min. Raw output data from the MS was converted to mzXML format using inhouse-developed software, and quantification of metabolites were performed by using the Metabolomics Analysis and Visualization Engine (MAVEN 2011.6.17, http://genomics-pubs.princeton.edu/mzroll/index.php) software suite. Peaks were matched to known standards for identification.

### Purification of genomic DNA

Genomic DNA from *C. crescentus* was purified according to the Puregene® DNA Handbook (Qiagen) protocol for gram negative bacteria. Cell lysis, RNA degradation, protein precipitation, and DNA precipitation were all performed as directed. DNA was left at room temperature for 3 days with gentle shaking to fully dissolve.

### qPCR to determine the ratio of oriC/ter

Hydrolysis probe qPCR was performed on purified genomic DNA from synchronized cells(43). Primer sequences for the *oriC* and *ter* regions are available upon request. Internal probes had 5’ fluorescein reporters and 3’ TAMRA quenchers. qPCR was performed with PrimeTime Gene Expression Master Mix (IDT), and 20µL reactions were prepared according to manufacturer’s directions in a MicroAmp optical 96 well plate. Genomic DNA was diluted 1:100. Reactions were conducted in a Quant Studio 7 Flex instrument with the following thermocycler program: 95°C for 3 min and 40 cycles of 95°C for 15 sec and 60°C for 60 sec. The average C_T_ value for technical replicates was used to calculate relative copy number of *oriC* with the ΔΔC_T_ method.

### High-throughput sequencing to determine replication rates

Genomic DNA was purified from synchronized NA1000 strains. Illumina sequencing libraries were prepared using the tagmentation-based and PCR-based Illumina DNA Prep kit and custom IDT 10bp unique dual indices (UDI) with a target insert size of 320bp. Sequencing was performed on an Illumina NovaSeq 6000, producing 2x151bp paired-end reads. Demultiplexing, quality control, and adapter trimming was performed with bcl-convert (v4.1.5).

2.67M reads were collected per sample. Reads were mapped to the NA1000 genome with bowtie2 (v2.3.5.1) and sorted with samtools (v1.10). The number of reads per nucleotide position was determined with bedtools (v2.27.1). Read counts were averaged over 5,000bp windows and plotted as a function of chromosome position.

## Data availability

Illumina sequencing data was uploaded to the Sequence Read Archive under BioProject PRJNA1096337.

## Acknowledgements

We thank Jin Yang and Jue D. Wang for providing the nucleotide extraction protocol and ppGpp substrates. We thank Rachel Salemi for assistance with time-lapse microscopy and image analysis. This work was funded by the National Institutes of Health (NIH) grant 1R35GM150652 and the National Science Foundation (NSF) grant 1715710 to D.A.N. The funders had no role in study design, data collection/interpretation, or the decision to submit the work for publication.

